# Risk-Window Palbociclib Exposure Delays ErbB2-Driven Mammary Tumorigenesis and Suppresses Mammary Epithelial Cell Stemness

**DOI:** 10.64898/2026.09.14.751306

**Authors:** Yongxuan Liu, Amanda B. Parris, Zhikun Ma, Limin Wei, Yijun Qi, Xiaohe Yang

**Author notes:** Correspondence: Xiaohe Yang Ph.D. or Yijun Qi Ph.D. These authors contributed equally to this work (Yongxuan Liu, Amanda B. Parris).

## Abstract

ErbB2 overexpression occurs in 15–20% of invasive breast cancer (BC) and inactivation of Cyclin D1-CDK4/6 axis reduce mammary stem/progenitor cells in ErbB2-driven tumorigenesis. Here, we explored the preventive role of short-term palbociclib intervention in MMTV-ErbB2 mice. Palbociclib significantly inhibited the proliferation and stemness of ErbB2-overexpressed BC cells in vitro and in vivo. Furthermore, short-term palbociclib exposure during the early premalignant risk window significantly delayed mammary tumor development and reduced tumor multiplicity, accompanied by the suppression of epithelial proliferation and ductal/alveolar morphogenesis in premalignant tissues, as well as prolonged tumor-free survival compared with controls. Importantly, palbociclib reduced the luminal epithelial (CD24^high^/CD49f^low^), mammary reconstitution unit-enriched (CD24^high^/CD49f^high^) subpopulations, and luminal progenitor/TIC-enriched (CD61^high^/CD49f^mid^) subpopulations. These changes were accompanied by diminished mammary epithelial cell stemness functions, including colony-forming, mammosphere-forming, and 3D growth activities of mammary epithelial cells. Mechanistically, palbociclib-treated tissues showed inhibition of the Cyclin D1-CDK4/6-RB-E2F axis and coordinated attenuation of ER-, ErbB2- and Wnt/β-catenin-associated signaling. Together, we demonstrate that short-term CDK4/6 inhibition during a premalignant risk window produces a sustained delay in ErbB2-driven mammary tumorigenesis, associated with remodeling of the mammary epithelial hierarchy and suppression of proliferative and stem/progenitor-associated activity, suggesting CDK4/6 inhibition as a strategy for ErbB2-positive BC prevention.

**Graphic Abstract:** 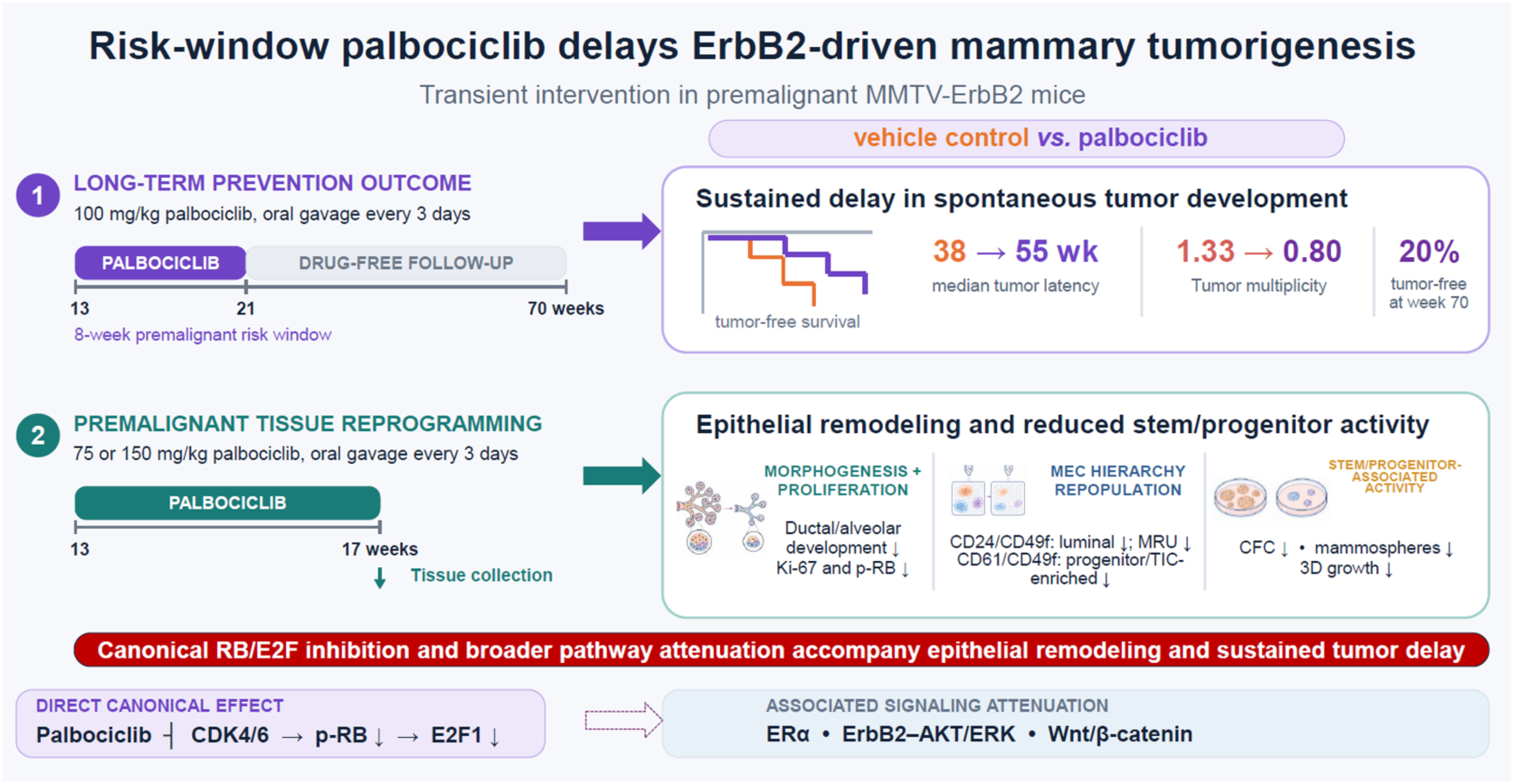

## 1. Introduction

With more than 2.3 million new cases and 685,000 deaths in 2020, the projected burden of breast cancer (BC) is predicted to increase to over 3.0 million new cases and 1.0 million deaths in 2040[1,2]. Although substantial advances in diagnosis and treatment have improved the clinical prognosis of BC, it is still a major challenge to reduce the burden of BC, highlighting the critical importance of effective prevention, particularly in high-risk populations. Human epidermal growth factor receptor (ErbB2/HER2) overexpression can be found in 15–20% of invasive BC, which is associated with an aggressive phenotype and poor clinical outcome [3–6]. Despite that the continuous development of ErbB2-targeted therapies has substantially improved the outcome of patients with ErbB2-overexpressed BC, therapeutic resistance against ErbB2 targets restrains their long-term efficacy [6]. Therefore, development of an effective preventive strategy for high-risk populations are urgently needed to reduce the burden of ErbB2-positive BC.

Mammary tumorigenesis is increasingly recognized to be a multistep process initiated from mammary precursor lesions, eventually transforming into an overt malignant state. The mammary gland is organized as a hierarchical system that comprises mammary stem cells (MaSCs), progenitor cells, and differentiated mammary epithelial cells (MECs), and disruption of this hierarchy gives rise to tumor initiation. MaSCs/progenitor cells and tumor-initiating cells (TICs) possessing the capacities of enhanced self-renewal and proliferation have been implicated in development and progression of BC [7]. Increasing evidence has underscored the close relationship between BC risk and MaSC/TIC stemness [8,9]. Thus, targeting MaSCs during the premalignant stages may represent a viable and effective strategy for BC prevention.

Accumulating evidence indicates that targeting Cyclin D1-Cyclin-dependent kinase 4/6 (CDK4/6) axis has the potential to abrogate mammary stem/progenitor cells in the context of ErbB2-driven tumorigenesis [10–12]. Several studies using CyclinD1^KE/KE^ mice (generated by the point mutant Cyclin D1-K112E knockin) demonstrated that Cyclin D1-associated kinase activity is required for the self-renewal, regeneration, and luminal lineage differentiation of mammary progenitor cells [13]. Loss of Cyclin D1 kinase activity impaired the ability of maintenance of progenitor populations, which served as targets for MMTV-ErbB2-mediated transformation, and protected mice from ErbB2-driven mammary tumorigenesis [10–14]. These findings provide a strong rationale to reduce the risk of BC incidence by specially targeting the Cyclin D1-CDK4/6 axis. Palbociclib, a selective CDK4/6 inhibitor, may have the potential to suppress mammary stem/progenitor cell function and tumor-initiating capacity in ErbB2-driven premalignant lesions.

In this study, we investigated whether short-term exposure to palbociclib during early risk window can suppress ErbB2-driven mammary tumorigenesis through modulation of mammary epithelial hierarchy and stem/progenitor cell function. By using ErbB2-positive BC cell lines and MMTV-ErbB2 transgenic mice, we examined the effects of palbociclib on cell proliferation, mammary stem/progenitor-associated phenotypes, composition of MECs, and tumor initiation. Our findings demonstrate that short-term intervention by targeting CDK4/6 can suppresses mammary cell stemness, remodel the composition of MEC in premalignant lesions, as well as significantly defer mammary tumorigenesis, providing a novel avenue for prevention of BC during early risk-window.

## 2. Materials and methods

### 2.1 Antibodies and Reagents

Palbociclib was purchased from LC Laboratories (Woburn, MA, USA). For in vitro experiments, palbociclib was dissolved in sterile water as a 10 mM stock solution and stored at −20 °C until use. For in vivo administration, palbociclib was suspended in sodium lactate buffer (50 mM, pH 4.0) containing 0.5% methylcellulose 400, which served as the vehicle for all animal experiments [15]. Primary antibodies including p-p70S6K, p70S6K, p-4EBP1, 4EBP1, p-ERBB2, p-AKT, p-ERK1/2, p-ER, active-β-catenin, β-catenin, p-Rb, Rb, c-MYC, CDK1, DVL2, c-JUN, LRP6, OCT4, p-ERα/118, p-ERα/167, c-MYC, and horseradish peroxidase-labeled goat anti-rabbit/mouse secondary antibodies were purchased from Cell Signaling (Danvers, MA). Antibodies including BCL-2, CDK4, E2F1, ERK2, ERα, Cyclin D1, AKT, Actin, KLF4, and NANOG, were obtained from Santa Cruz Biotechnology (Santa Cruz, CA). Antibodies of ERBB-2, SOX2 were purchased from EMD Millipore (Billerica, CA).

### 2.2 Cell Culture

BC cell lines of SKBR-3 (ATCC, Cat# HTB-30, RRID: CVCL_0033) and BT-474 (ATCC, Cat# HTB-20, RRID: CVCL_0179) were purchased from the American Type Culture Collection (ATCC, Manassas, VA, USA) and maintained in DMEM/F-12 culture medium supplemented with 10% fetal bovine serum, penicillin (100 U/mL), and streptomycin (100 µg/mL) in a humidified incubator with 5% CO2 at 37 ℃. Murine mammary tumor cell line 85819 was established from spontaneous mammary tumors of MMTV-ErbB2 transgenic mice as previously described [7]. Cell treatments were specified in the corresponding experiments.

### 2.3 Cell Proliferation Assay

Cells were seeded into 96-well plates at 800 cells/well and cultured for 24 h before treatment. The cells were then treated with palbociclib at the indicated concentrations for 5 days, followed by incubation with Cell Counting Kit-8 (CCK-8) for 4 h. Absorbance was measured at 450 nm using a microplate reader (BioTek; Winooski, VT, USA). Viable cell fraction of each group, based on 4 parallel samples, was calculated relative to the controls, which were normalized to 100% survival.

### 2.4 Clonogenic Assay

Cells were seeded into 6-well plates at a density of 600 cells per well and treated with palbociclib at the indicated concentrations for 14 days. Colonies were then fixed with methanol and stained with 0.5% crystal violet. Colony images were acquired using a Nikon SMZ745T microscope. Colonies were quantified using ImageJ software. Data based on triplicates were analyzed using GraphPad Prism software.

### 2.5 Cell Cycle Analysis

SKBR-3 and BT-474 cells were treated with indicated concentrations of palbociclib for 24 h. Then, the cells were trypsinized and fixed in ice-cold 70% ethanol (added dropwise) at −20 °C overnight. Fixed cells were centrifuged to form a cell pellet and the supernatant was removed. The cell pellets were resuspended and incubated in PBS containing 0.2% triton X-100, 500 μg/ml RNase A, and 33 μg/ml propidium iodide at 37 °C for 45 min. Cells in different cell cycle phases were analyzed using a Guava EasyCyte 8 flow cytometer (Millipore; Billerica, MA, USA). The percentages of the cells in each phase of cell cycle were analyzed with ModFit software [16]. Representative data from three sets of repeats were presented.

### 2.6 ALDEFLUOR assay

The ALDH1 activity was evaluated using ALDEFLUOR assay kit (STEMCELL Technologies, Vancouver, Canada), SKBR-3 cells were treated with palbociclib at the indicated concentrations for 24 h and harvested. The cells were suspended in ALDEFLUOR assay buffer at 3 × 10⁵ cells/mL and incubated with the activated ALDEFLUOR substrate according to manufacturer’s instructions. Parallel negative controls were prepared by treating a subset of cells with the ALDH1 inhibitor DEAB prior to substrate addition. Following incubation at 37 °C for 45 min, the cells were washed and resuspended in cold buffer. The ALDEFLUOR-positive population was defined based on the DEAB control, and ALDH1 activity in viable cells identified by 7-AAD exclusion was analyzed using a Guava EasyCyte 8 flow cytometer (Millipore; Billerica, MA, USA). The percentage of ALDH-positive cells from triplicate samples was quantified and statistically analyzed.

### 2.7 Syngeneic Tumor Model

Murine 85819 cells (1 × 10⁶ cells in a 100 μL suspension in a 1:1 mixture of serum-free medium and Matrigel) were injected subcutaneously into the right flank of 8-week-old female FVB/N recipient mice which were purchased from The Jackson Laboratory (Bar Harbor, ME, USA). Mice were randomized into three groups (N = 8) on day 4. Mice received either vehicle control, 75 mg/kg palbociclib, or 150 mg/kg palbociclib by oral gavage every 3 days from day 4 till the termination of this experiment. Tumor growth was monitored by caliper measurement twice a week. Tumor volume was calculated using the formula: Volume = longest diameter × shortest diameter^2^ × 0.5. At the endpoint, tumors were harvested and fixed in 10% neutral buffered formalin for immunohistochemistry (IHC).

### 2.8 Animals and Treatments

Female FVB/N-MMTV-ErbB2 transgenic mice were purchased from the Jackson Laboratory (Bar Harbor, ME, USA) and maintained on an estrogen-free AIN-93G diet. All animal procedures were approved by the Institutional Animal Care and Use Committee.

For tumor development assessment, 13-week-old mice (n = 15 per group) were randomized to two groups, including vehicle control group or palbociclib treatment group (100 mg/kg) by oral gavage every 3 days for 8 weeks. Spontaneous mammary tumor development was monitored twice weekly by palpation and caliper measurement. Tumor latency (the time from birth to detection of the first palpable mammary tumor), cumulative tumor incidence and tumor-free survival were analyzed using Kaplan-Meier survival curves. Differences in survival outcomes between the groups were evaluated using the log-rank test. Tumor multiplicity was defined as the total number of mammary tumors developed per mouse during the observation period. Statistical significance was determined using a two-tailed Mann-Whitney test.

For the evaluation of palbociclib-treated premalignant mammary tissues, mice were treated with vehicle or palbociclib at 75 or 150 mg/kg every 3 days from 13 to 17 weeks of age. Mammary tissues were collected from 3 mice per group for analyses of whole-mount, IHC, flow cytometry, and protein expression.

### 2.9 Western Blot

Total proteins were extracted from cells or homogenized mammary tissues as described in our previous report^3,4^. Protein concentration was quantified using a BCA Protein Assay Kit (Thermo Scientific Pierce). Equal amounts of protein (30 µg/lane) were separated by electrophoresis on 10% or 12% SDS-PAGE gels and then transferred onto nitrocellulose membranes. Membranes were blocked in 5% milk for 2 h at room temperature and then incubated in primary antibodies (1:1,000 or 1:2,000 dilution) at 4 °C overnight. After washing, the membranes were incubated in secondary horseradish peroxidase-labeled antibodies for 1.5 h at room temperature. Protein bands were visualized using enhanced chemiluminescence (ECL) reagents (Thermo Fisher Scientific) and captured with an Azure imaging system.

### 2.10 Mammary epithelial cell isolation and flow cytometry analysis

Primary mammary epithelial cells were isolated from inguinal mammary glands using the protocol provided by a Gentle Collagenase/Hyaluronidase kit (Cat. No. 07919, STEMCELL Technologies). The glands were minced with a tissue chopper and digested with gentle collagenase/hyaluronidase at 37 ℃ for 2 h under constant rotation at 90 rpm. The resulting organoids were further dissociated with trypsin-EDTA and dispase (5 mg/mL)/DNase I (STEMCELL Technologies). The cell suspension was filtered using a 40 μm cell strainer.

For flow cytometry analysis of MEC populations, cells were blocked with CD16/CD32 to prevent non-specific binding and then stained with a panel of lineage markers (CD45, CD31, and Ter119) to exclude hematopoietic, endothelial, and erythroid cells, along with 7-AAD to identify viable cells. Cells were simultaneously stained with either CD61/CD49f or CD24/CD49f antibodies. Staining and gating strategies for lineage-negative (Lin−) cells and individual subpopulations were performed in accordance with previous publications [18–21].

### 2.11 Mammary gland whole-mount analysis

Inguinal mammary glands were excised from all involved mice, spread flat on glass slides, and fixed in Carnoy’s fixative (6:3:1 ratio of 100% ethanol: chloroform: glacial acetic acid) overnight at room temperature. Fixed tissues were washed in 70% ethanol, rehydrated, and stained with carmine alum overnight to visualize the epithelial structure, followed by sequential dehydration, clearing in xylene, and mounting with Permount (Thermo Fisher Scientific). Whole mounts were imaged and analyzed using a Nikon stereo microscope system (Nikon SMZ745T) to evaluate ductal growth, branching, and alveolar development.

### 2.12 Tumorsphere, mammosphere, and 3D culture assays

For tumorsphere assays, single-cell suspensions of BT-474 cells were seeded at 2,000 cells/well in ultra-low attachment 24-well plates containing EpiCult-B Human Medium (STEMCELL Technologies, Vancouver, BC, Canada), supplemented with 1× B27, 20 ng/mL EGF, 10 μg/mL insulin, 1 μg/mL hydrocortisone, 20 ng/mL bFGF, and 4 μg/mL heparin. The cells were incubated at 37 ℃ with 5% CO₂ for 7 days to assess primary sphere formation. The images of spheres with a diameter greater than 30 μm were counted and acquired. Primary spheres were then dissociated into single-cell suspensions and inoculated into new plates at a density of 1 × 10^3^ cells/well under identical conditions to assess the secondary sphere-forming efficiency. Both primary and secondary sphere assays were performed in triplicate.

For mammosphere assays, MECs isolated from the mammary glands as described above were seeded into ultra-low attachment 24-well plates at 2.5 × 10⁴ cells/well in EpiCult-B Mouse Medium (STEMCELL Technologies, Vancouver, BC, Canada) with the supplements. Culture and quantification of primary and secondary mammospheres were performed as described above for the tumorsphere assay.

For the 3D culture assay, primary MECs isolated from all involved mice were seeded into 48-well plates at 1.5 ×10⁴ cells/well in a chilled 1:1 mixture of growth factor-reduced Matrigel and EpiCult-B Mouse Medium and cultured for 10 days. The images of colonies were then stained with crystal violet, quantified, and acquired. Data from triplicate samples were statistically analyzed.

### 2.13 Colony-Forming cell assay

Primary MECs isolated as described above were seeded into 60-mm dishes at 4 × 10³ cells/plate. Cells were cultured in EpiCult-B Mouse Medium supplemented with 1 μg/mL hydrocortisone and 4 μg/mL heparin for 10 days. Colonies were then fixed with methanol and stained with Wright’s Giemsa stain. Colony images were acquired using Nikon SMZ745T microscope and imaging system. These assays were performed in triplicate.

### 2.14 Immunohistochemistry

Formalin-fixed, paraffin-embedded mammary tissues were sectioned and subjected to IHC staining to evaluate the expression of Ki67, phospho-RB, and Cyclin D1. Antigen unmasking was performed by heating sections in a citrate buffer (pH 6.0). Following primary antibody incubation, signal visualization was carried out using the avidin-biotin complex method (ABC-DAB system, Vector Laboratories). Sections were counterstained with hematoxylin, and brightfield images were captured using a Nikon Eclipse 80i microscope and Nikon Elements Imaging System Software.

### 2.15 Statistical analysis

Data are presented as mean ± standard error (SEM). Statistical analyses were performed using GraphPad Prism 10. Comparisons between two groups were conducted using Student’s t-test, and multiple-group comparisons were analyzed by one-way ANOVA followed by appropriate post hoc testing. Tumor-free survival was analyzed using the Kaplan-Meier method with log-rank testing. A p value < 0.05 was considered statistically significant.

## 3. Results

### 3.1 Palbociclib inhibits the proliferation of ErbB2 overexpressing BC cells

To evaluate the effect of palbociclib on ErbB2-overexpressing BC cells, SKBR-3 and BT-474 cells were treated with increasing concentrations of this drug. In both BC cell lines, cell viability assays showed significant dose-dependent inhibitions of cell proliferation, with IC₅₀ values of 2.94 μM and 0.55 μM for SKBR-3 and BT-474 cells, respectively (Figure 1A). Consistent with this, clonogenic assays also demonstrated marked reductions in both colony number and colony size following palbociclib treatment (Figure 1B). Cell cycle analysis revealed that palbociclib induced G0/G1 cell cycle arrest in SKBR-3 and BT-474 BC cells. In both cell lines, palbociclib increased the percentages of cells in G0/G1 phase accompanied by reductions in the percentages of cells in S phase, especially in the BT-474 cells, which is indicative of diminished proliferating cells (Figure 1C). These data indicate that palbociclib exerts an anti-tumor activity.

**Figure 1.**
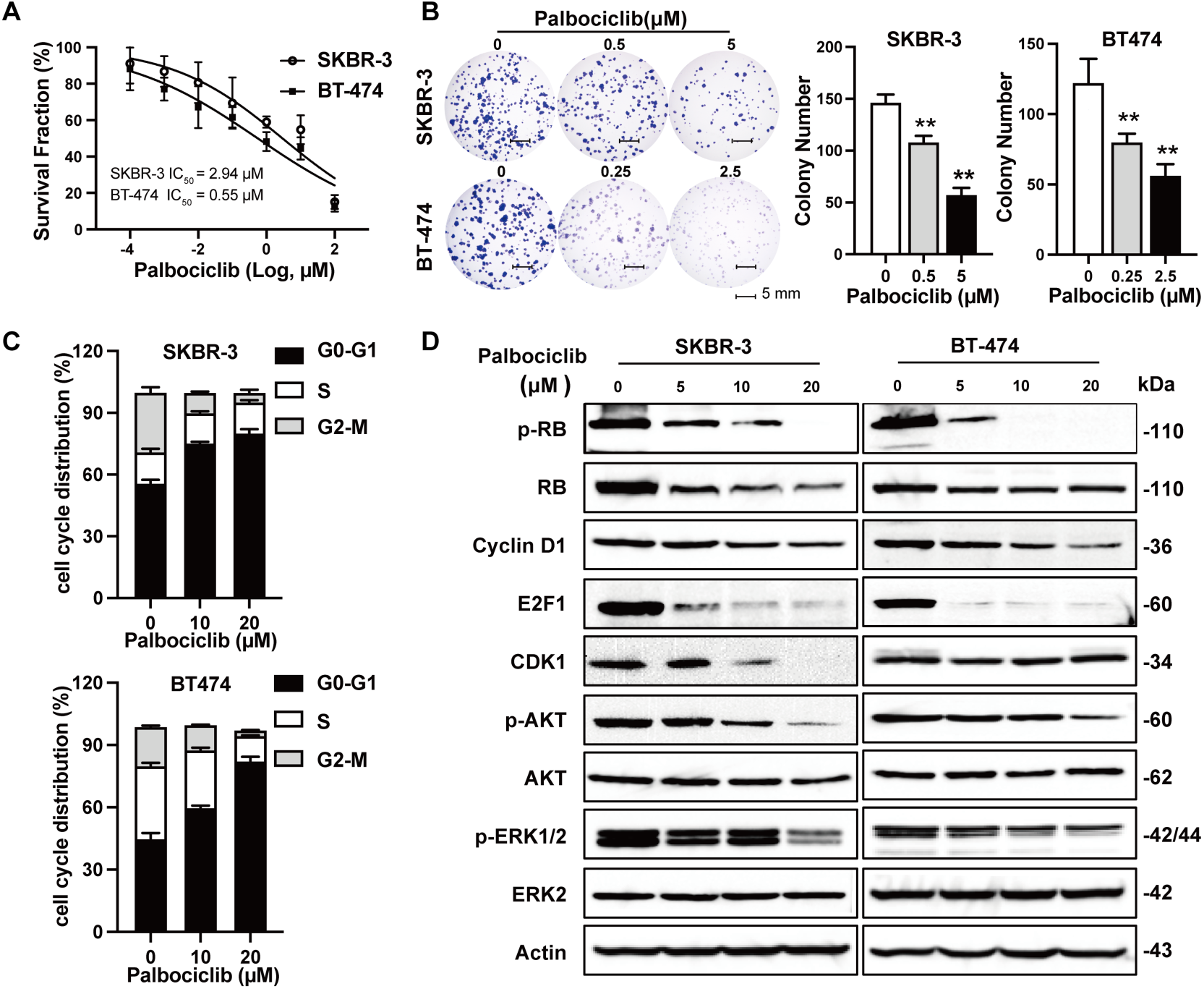
Palbociclib induces anti-proliferative effects in ErbB-2 overexpressing SKBR-3 and BT-474 cells line. **A)** CCK-8 assay was performed to compare cell viability in SKBR-3 and BT-474 cells treated with palbociclib (10^-4^, 10^-3^, 10^-2^, 10^-1^,1,10, 10^2^ μM) for 5 days. The average viable cell fraction for each sample (N = 4) is mean ± standard error (SEM). **B)** SKBR-3 (0, 0.5, 5**μ**M) and BT-474 (0, 0.25, 2.5**μ**M) were treated with palbociclib in triplicate for 14 days as part of the clonogenic assay. Then the colonies were fixed and stained with crystal violet. The average number of colonies that formed after 14 days is graphed. Representative images of crystal violet-stained colonies are shown in the right panel. Values are graphed as the mean ± SEM (\*\**p* < 0.01, as compared to the corresponding untreated control samples). **C)** SKBR3 and BT474 cells were treated with palbociclib (0, 10, 20 μM) for 24 h and then cell cycle progression was measured using fluorescence activated cell sorting analysis (N = 3). The average percentage of cells in G2/M, S, and G0/G1 phases of the cell cycle are graphed for SKBR-3 and BT-474 cells **D)** Palbociclib regulates signaling pathway that promote cell proliferation in ErbB-2 overexpressing breast cancer cells line. SKBR-3 and BT-474 cells were treated with palbociclib (0, 5, 10, 20 μM) for 24h, followed by western blot analysis of indicated markers in the cell cycle pathway and the Receptor Tyrosine Kinase (RTK) pathways

To investigate the underlying mechanisms, we examined cell-cycle and growth-associated signaling pathways that are mechanistically related to the effect of palbociclib. As expected, proliferation-associated proteins, including p-RB, E2F1, Cyclin D1, and CDK1, were significantly down-regulated by palbociclib in both BC cell lines (Figure 1D), consistent with the inhibition of CDK4/6-RB-E2F axis. In addition, the phosphorylation levels of AKT and ERK2 proteins were also remarkably diminished following palbociclib treatment. Together, these findings demonstrate that palbociclib effectively suppresses cell proliferation in ErbB2-overexpressing BC cells, which support our further studies in MMTV-ErbB2 mouse model in vivo.

### 3.2 Palbociclib inhibits the tumor stemness of ErbB2-overexpressing BC cells

Because cancer stem/progenitor cells play a critical role in tumor initiation and therapeutic resistance [22–24], we next explored whether palbociclib impairs the tumor stemness of BT-474 and SKBR-3 cells. Tumorsphere formation assay revealed that palbociclib significantly reduced the capacity of tumorsphere formation in both the primary and secondary cell culture using ultra-low attachment plates (Figure 2A). Flow cytometric analysis using an ALDEFLUOR assay kit showed that ALDH-positive cells were significantly reduced by palbociclib compared with untreated controls (Figure 2B), indicating suppression of the tumor stemness in SKBR-3 cells as well. In line with these stemness-related phenotypes, palbociclib treatment also resulted in the markedly down-regulation of stemness-associated proteins, including KLF4, NANOG, and SOX2 in both SKBR-3 and BT-474 cells (Fig. 2C). Together, these findings demonstrate that palbociclib has the ability to impair the cancer stemness of ErbB2-overexpressing BC cells, thus leading to low potential of proliferation.

**Figure 2.**
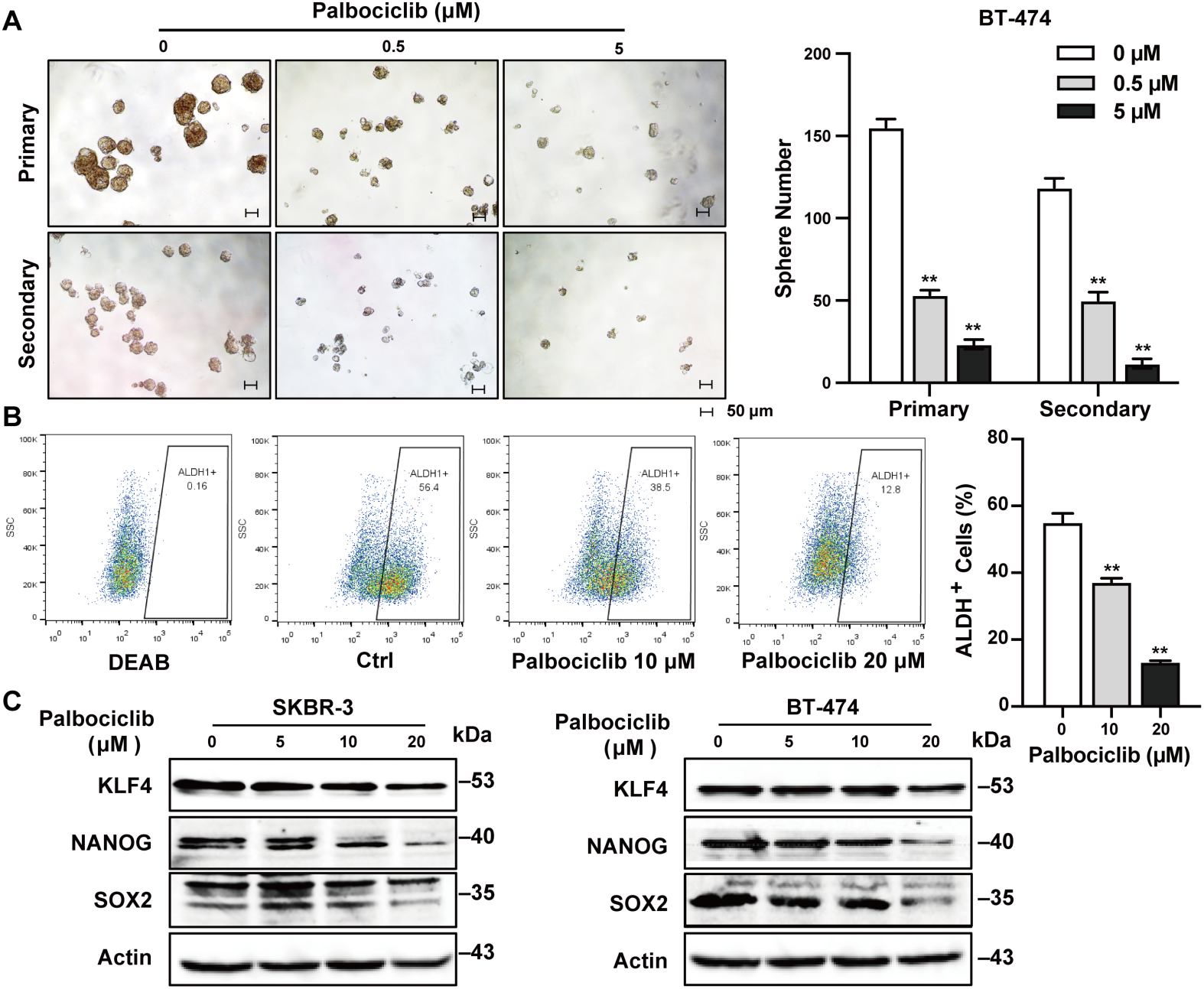
Palbociclib inhibits the stemness of ErbB-2 overexpressing breast cancer in vitro. **A)** BT-474 cells were cultured in vitro and subjected to primary and secondary tumor sphere assays. In the primary tumor sphere assay, cells were initially treated with palbociclib (0, 0.5, or 5 μM) for 7 days for primary sphere formation. Then primary tumor spheres were harvested and replated for another 7 days under identical incubation conditions to form secondary spheres. Primary and secondary tumor sphere formations were recorded. Values are presented as the means ± SEM (\*\**p* < 0.01 as compared to the corresponding untreated control samples). **B)** SKBR-3 cells were treated with palbociclib (0, 10, or 20 μM) for 24 h, followed by quantification of ALDH-positive cells. The percentage of ALDH-positive cells was determined using the ALDEFLUOR detection kit with flow cytometry. Values are presented as the means ± SEM (\*\**p* < 0.01 as compared to the control for each treated group). **C)** Palbociclib, at the indicated concentrations, for 24 h, followed by Western blot analysis of stemness markers.

### 3.3 Palbociclib inhibits exogenous ErbB2-driven cell proliferation in vitro and syngeneic tumor growth in vivo

Because BC cell lines SKBR-3 and BT-474 have upregulated expression of endogenous ErbB2, we then evaluated the effect of palbociclib in an exogenously ErbB2-overexpressing cell line 85819. In vitro, palbociclib consistently inhibited the abilities of cell proliferation and colony formation in a dose-dependent manner in a mammary tumor cell line 85819 derived from MMTV-ErbB2 transgenic mice (Figure 3A), indicating the sensitivity of this in vitro transgenic cell model to CDK4/6 inhibition. We next assessed the in vivo efficacy of palbociclib. In the syngeneic tumor model, mice implanted subcutaneously with 85819 cells were treated with palbociclib (75 or 150 mg/kg) every 3 days for a total of 29 days, starting on day 4 after subcutaneous inoculation (Figure 3B). Both dosages of palbociclib significantly suppressed the subcutaneous tumor growth compared with the vehicle control group. Specifically, the high dosage yielded a 61.47% decrease in tumor weight in comparison with a 48.38% reduction using the low dosage, indicating a dose-dependent inhibition in tumor growth (Figure 3B), In addition, there was no significant difference in mice weight, suggesting that both dosages were well tolerated in our animal experiment. Consistent with these findings, IHC analysis of grafted tumors showed significantly decreased protein levels of Ki67, p-RB and SOX2 in palbociclib-treated tumors (Figure 3C). Together, these results demonstrate that palbociclib can effectively inhibit the growth of ErbB2-driven mammary tumors in vitro and in vivo, supporting further evaluation of the activity of palbociclib in premalignant mammary tissues in MMTV-ErbB2 transgenic mice.

**Figure 3.**
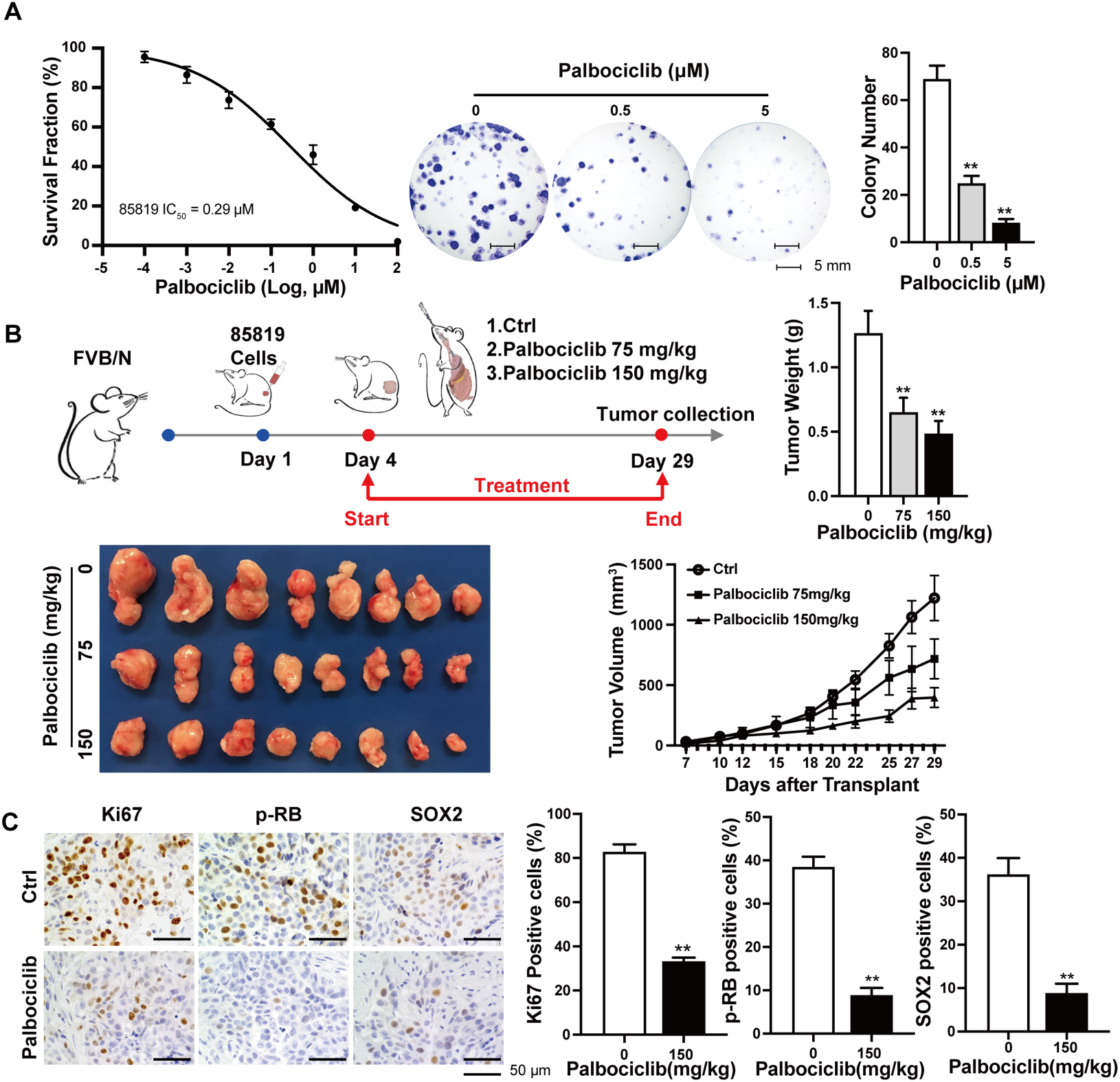
Palbociclib inhibits cell proliferation of 85819 cells in vitro and syngeneic grafted tumors. **A)** CCK-8 assay was performed to cell viability in 85819 cells treated with palbociclib (10-4, 10-3, 10-2, 10-1,1,10, 102 μM) for 5 days. The average viable cell fraction for each sample (N = 4) is mean ± SEM. 85819 (0, 0.5, 5 μM) were treated with palbociclib in triplicate for 14 days as part of the clonogenic assay. Then the colonies were fixed and stained with crystal violet. The average number of colonies that formed after 14 days is graphed. Representative images of crystal violet-stained colonies are shown in the right panel. Values are graphed as the mean ± SEM (\*\**p* < 0.01, as compared to the corresponding untreated control samples). **B)** FVB/N mice were injected with 85819 cells and then treated with palbociclib (75 mg/kg and 150 mg/kg every 3 days) for 29 days. Palpable tumor sizes were recorded twice a week. Harvested tumors from animals with different treatments. Average tumor weights in the control and palbociclib treatment groups are presented as mean ± SEM Tumor growth curve of different treatment groups. **C)** Ki67, p-RB and SOX2 IHC staining of proliferating and stemness MECs in control and treated tumors. Values are presented as the mean ± SEM (\*\**p* < 0.01).

### 3.4 Early and short-term intervention by palbociclib suppresses mammary tumorigenesis in MMTV-ErbB2 transgenic mice

Breast carcinogenesis involves a multistep and complicated process that gradually accumulates a number of genetic and epigenetic aberrations, including ErbB2. We next sought to investigate whether short-term palbociclib exposure during the premalignant risk window could suppress mammary tumorigenesis in MMTV-ErbB2 transgenic mice.

Mice were treated with palbociclib (100 mg/kg) every 3 days for 8 weeks during weeks 13–21, followed by closely monitoring the spontaneous tumor formation (Figure 4A). As shown in Figure 4B, palbociclib treatment significantly deferred the onset of mammary tumor, as well as markedly extended the tumor-free survival compared with the control group. The initial manifestation of palpable tumors started to appear at week 25 in the control mice versus at week 34 in the palbociclib group, respectively. Notably, three mice in the palbociclib group remained tumor-free till the termination of the animal experiment at week 70 in this study. Accordingly, the median tumor latency was significantly increased in mice treated with palbociclib compared with those of control mice (55 vs. 38 weeks, p < 0.01). Importantly, palbociclib also significantly decreased tumor multiplicity (0.80 ± 0.11 vs. 1.33 ± 0.16 tumors per mouse in the palbociclib and control groups, respectively), further supporting the long-term inhibitory effect of palbociclib on mammary tumorigenesis (Figure 4B).

**Figure 4.**
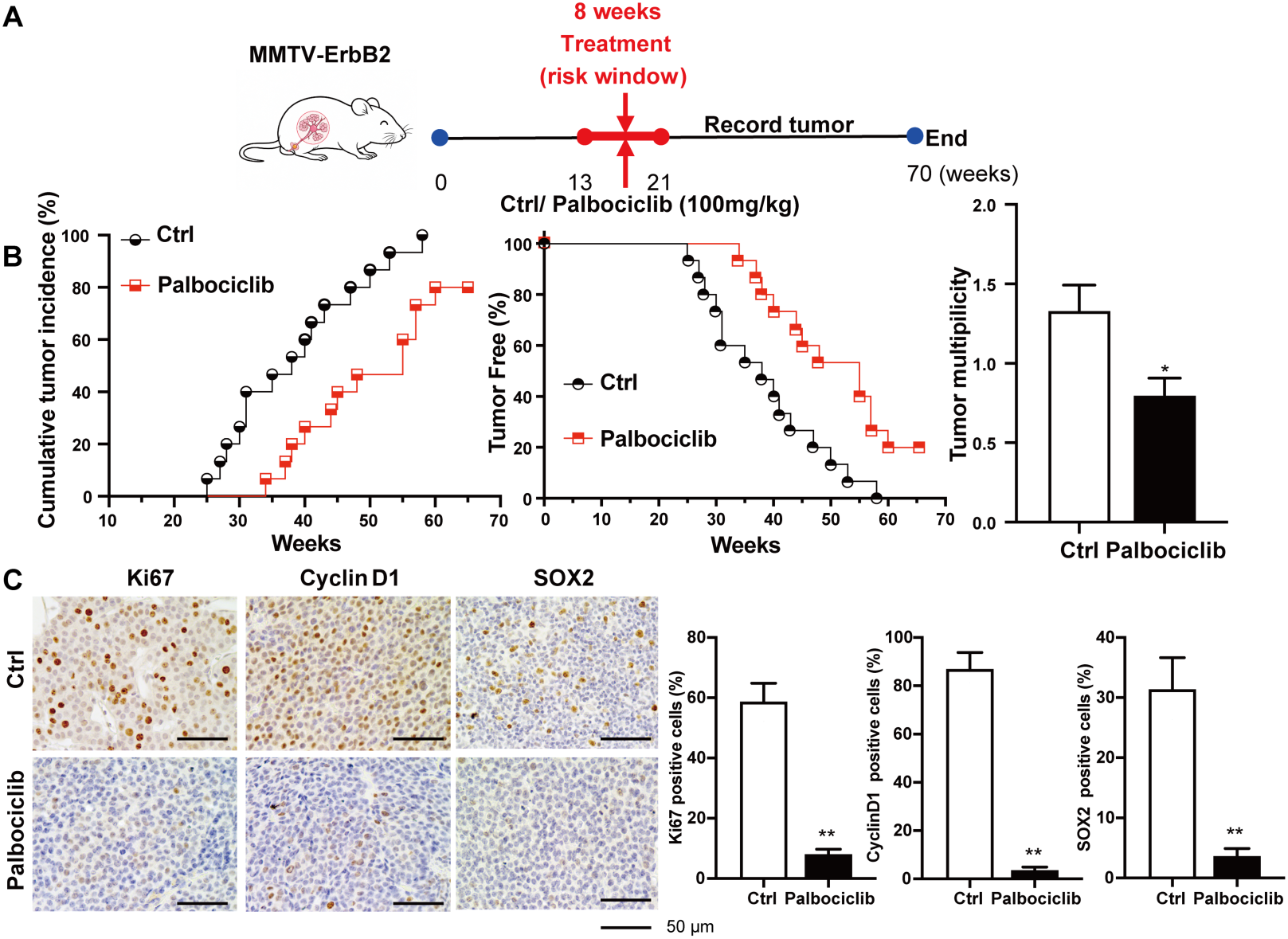
Short-term palbociclib exposure during the risk window increases mammary tumor latency in MMTV-ErbB2 transgenic mice. **A)** MMTV-ErbB2 transgenic mice were treated with control (n = 15) or palbociclib (n = 15) every 3 days from 13 weeks until 21 weeks of age. **B)** Tumor development was monitored twice a week from 20 to 70 weeks of age. The percentage of cumulative tumor incidence and tumor-free in control and palbociclib treated mice is shown in the Kaplan– Meier curves (*p* = 0.003). Tumor multiplicity in two mice groups was presented as mean ± SEM. Statistical significance was determined by Mann-Whitney test. \**p* < 0.05. **C)** IHC for Ki67, Cyclin D1 and SOX2 in control and palbociclib treated tumors. Values are presented as the means ± SEM (\*\**p* < 0.01).

To further determine whether early exposure to palbociclib during premalignant stages influenced the biological characteristics of tumors, we evaluated proliferation-associated markers by IHC. Tumors from palbociclib-treated mice exhibited significantly reduced staining of Ki67, Cyclin D1 and SOX2 compared with control tumors (Figure 4C), indicating decreased proliferative activity, likely due to persistent suppression of cell cycle progression signaling and stemness. Together, these findings demonstrate that short-time exposure to palbociclib during a premalignant risk window confers sustained protection against ErbB2-driven mammary tumor initiation and development.

### 3.5 Palbociclib inhibits mammary epithelial proliferation and morphogenesis in premalignant mammary tissues

Aberrant mammary epithelial cell proliferation, hyperplasia, and ductal morphogenesis precede the formation and development of malignant transformation in MMTV-ErbB2 transgenic mice and are associated with increased mammary tumor susceptibility [25,26]. To investigate the mechanisms underlying palbociclib-mediated tumor suppression, we examined its effects on mammary morphogenesis and epithelial cell proliferation during the premalignant stages. MMTV-ErbB2 transgenic mice were treated with vehicle or palbociclib (75 or 150 mg/kg) between 13 and 17 weeks of age, and mammary glands were analyzed afterwards. Whole-mount analysis revealed that palbociclib markedly suppressed mammary ductal development, evidenced by significant inhibition of epithelial cell density, lateral branching, and alveolar budding compared with those from control mice (Figure 5A). These changes indicate the impeded expansion of mammary epithelial cells and morphogenic development in premalignant tissues, which may account for the delayed onset of neoplasia in palbociclib-treated mice.

**Figure 5.**
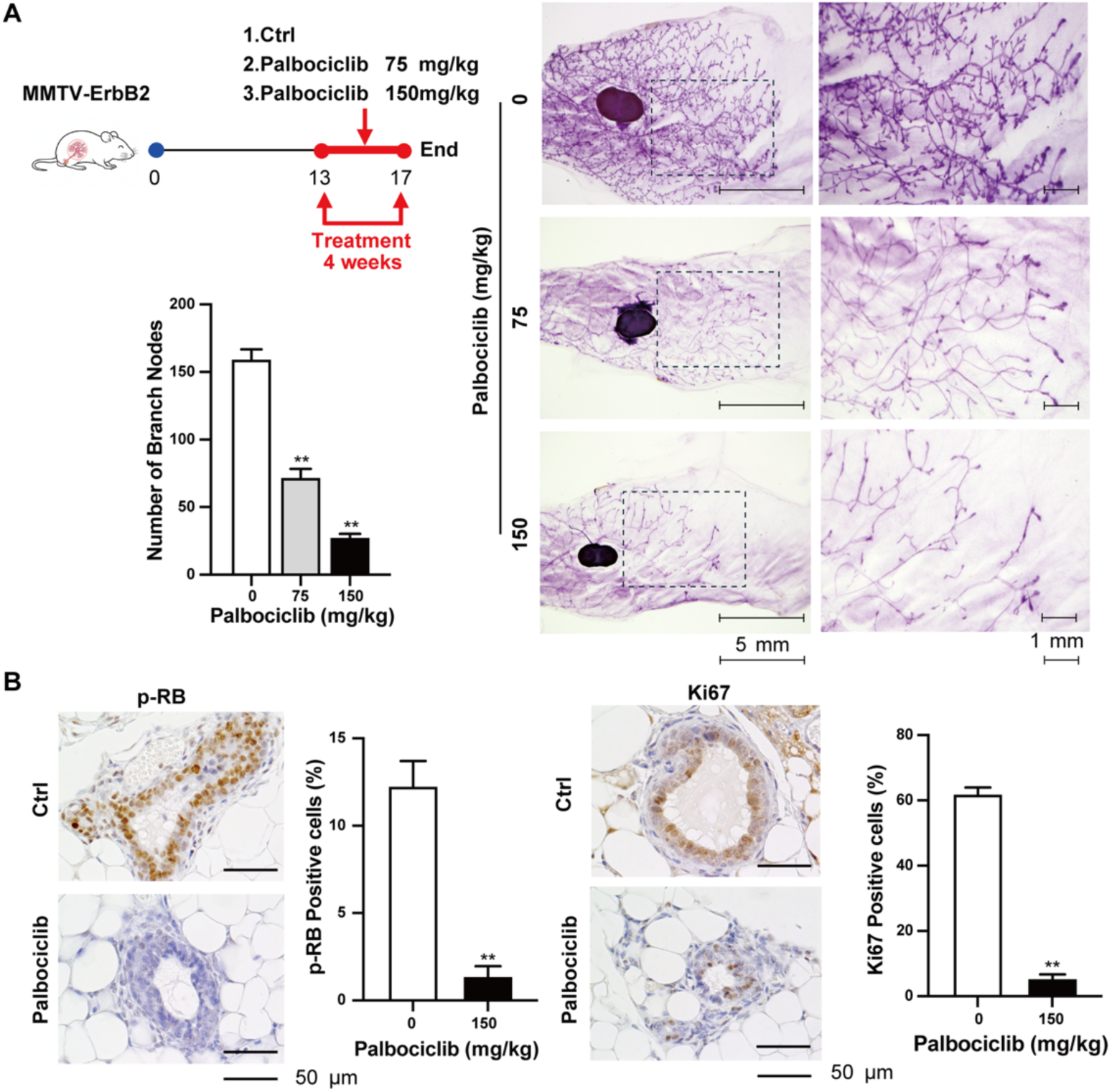
Palbociclib inhibits morphogenesis and proliferation of MECs in premalignant mammary tissues. **A)** Mammary glands were collected from 17-weeks-old mice in control and palbociclib treated groups (75 or 150 mg/kg every 3 days for 4 weeks). Representative images are shown at 2x and 4x magnification. The average number of side branches per field is graphed as the mean ± SEM in the light panel (\*\**p* < 0.01, as compared to the corresponding untreated control samples) **B)** IHC staining of phospho-RB (p-RB) and Ki67 in mammary tissues with different treatments. Brown staining in the mammary gland tissue sections represents positive mammary epithelial cells. Values are presented as the means ± SEM (\*\**p* < 0.01).

To determine the cellular basis of these morphological alterations, mammary tissues were analyzed for Ki67 and p-RB expression. Palbociclib treatment significantly reduced Ki67-positive epithelial cells in a dose-dependent manner, indicating suppressed epithelial cell proliferation (Figure 5B). In addition, palbociclib also reduced RB phosphorylation, confirming the inhibition of CDK4/6-RB signaling in vivo. Together, these findings demonstrate that palbociclib suppresses premalignant mammary epithelial cell expansion and morphogenesis, supporting an effective prevention of mammary tumorigenesis by targeting CDK4/6 during early risk-window.

### 3.6 Palbociclib reduces MEC subpopulations by reprogramming of MaSCs differentiation

Accumulating evidence indicates that MaSCs regulated by ErbB2 signaling, which are important in mammary gland development, are also capable of promoting carcinogenesis via oncogenic transformation into TICs [27–30]. We therefore investigated whether palbociclib exposure during the premalignant risk window alters MEC composition and stem/progenitor-associated subpopulations. MECs from MMTV-ErbB2 transgenic mice were first analyzed using CD24 and CD49f expression profiles, which distinguish among luminal cells, basal/myoepithelial cells (Myo), and mammary reconstitution units (MRUs). Luminal compartments are comprised of progenitor cells with proliferative properties, while MRU fractions are enriched with MaSCs. As shown in Figure 6A, palbociclib treatment significantly altered the pattern of distribution of MEC in premalignant mammary tissues. Palbociclib induced a statistically significant decrease in both the luminal and MRU subpopulations that are enriched with mammary stem cells. In contrast, there was a noticeable increase in Myo subpopulation although the difference was not statistically significant. These data demonstrate that palbociclib impairs the hierarchical composition of mammary gland.

**Figure 6.**
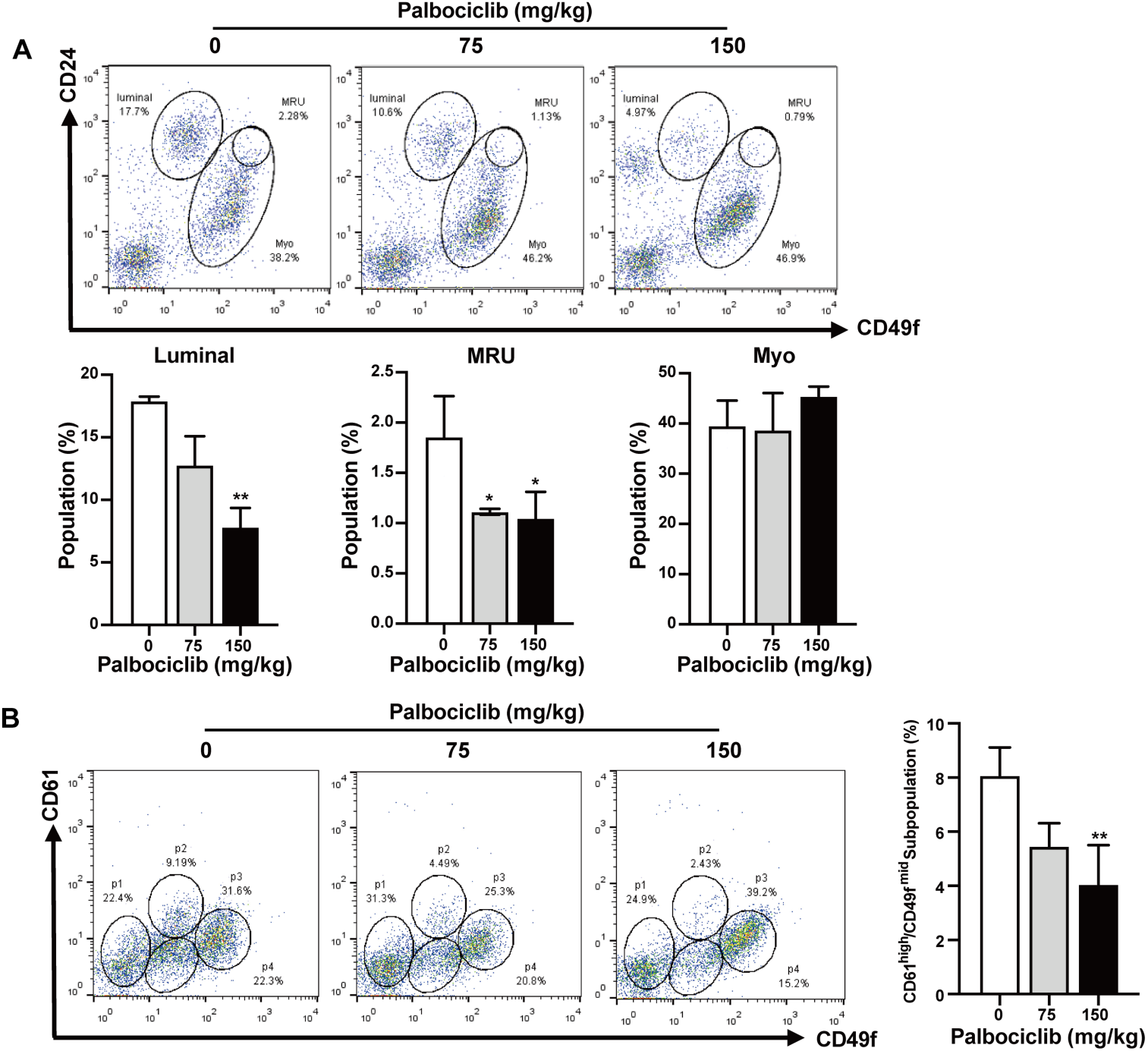
Palbociclib targets mammary stem cell populations and cancer stem cell/tumor-initiating cell populations in premalignant tissues from MMTV-ErbB2 mice. Primary MECs were isolated from 17-weeks-old mice that were gavaged the vehicle control or palbociclib (75 or 150 mg/kg every 3 days for 4 weeks). Cells were labeled with fluorescent antibodies for flow cytometry analysis of different MEC populations (N = 3). **A)** FACS analysis of MEC subpopulations based on CD24/CD49f profiles. Representative images of CD24/CD49f flow cytometry plots are shown. Percentage of cells from luminal, basal/myoepithelial (myo) and MRU (mammary reconstructive units) were analyzed. **B)** Isolated MECs were labeled with CD61/CD49f markers and analyzed with flow cytometry (N = 3). Representative images of CD61/CD49f flow cytometry plots are shown. The percentage of MECs of p2 (CD61highCD49fmid) subpopulation in mammary glands were compared between different treatments. Values are graphed as the mean ± SEM (\**p* < 0.05, \*\**p* < 0.01, as compared to the corresponding untreated control samples)

To characterize the specific cell populations involved in tumor initiation and its outcome as preventive targets, CD61/CD49f cell-surface markers were used to isolate these subpopulations from erbB-2 overexpressing mammary tumors [17,19]. In MMTV-ErbB2 transgenic mice, the CD61high/CD49fmid subset (p2) is enriched for TICs [19]. We found that palbociclib treatment significantly reduced the proportions of CD61high/CD49fmid cells compared with that of control mammary tissues (Figure 6B), indicating selective suppression of TICs that are enriched in MECs subpopulations.

### 3.7 Palbociclib suppresses the stemness of MEC in premalignant mammary tissues

Based on the above findings, we next sought to determine whether the palbociclib-induced changes in MEC subpopulations reflect attenuation of stemness of mammary stem/progenitor cells. To address this issue, we performed Colony-Forming cell (CFC), 3D culture and mammosphere assays using MECs isolated from the mammary glands of control and palbociclib-treated mice. CFC assays in Figure 7A showed that palbociclib significantly reduced the colony-forming ability, with the greatest inhibition observed in the high-dose group. Consistently, both mammosphere and 3D colony formation were markedly decreased following palbociclib treatment (Figure 7B, C). Palbociclib exposure significantly reduced SOX2-positive epithelial cells from premalignant mammary tissues, indicating attenuation of mammary epithelial stemness (Figure 7D). Correspondingly, the protein levels of the stemness-associated transcription factors NANOG, OCT4, KLF4, and SOX2 were dose-dependently reduced following palbociclib treatment (Figure 7E), supporting suppression of mammary stem/progenitor cell properties.

**Figure 7:**
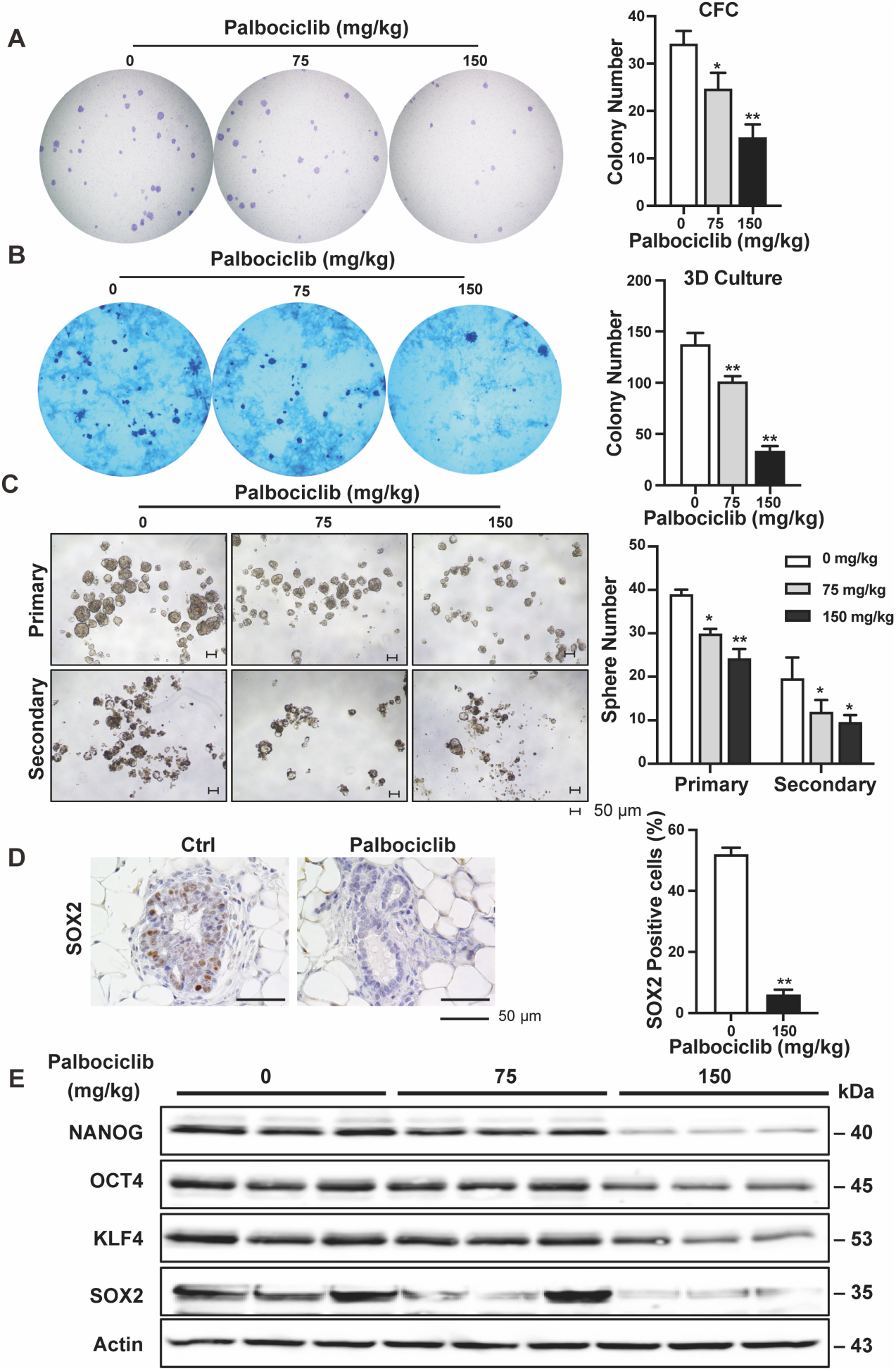
Palbociclib impairs the stemness of MECs form MMTV-ErbB2 mice. Primary MECs were harvested from 17-weeks-old mice that were treated with control or palbociclib (75 or 150 mg/kg every 3 days for 4 weeks). **A)** Primary MECs were plated for a CFC assay (N = 3). After 10 days, the cells were fixed and stained as previously described. The number of colonies in three mice groups was recorded. **B)** Colony formation of MECs derived from the three different treated mice group in 3D matrigel. After 10 days, Colonies were then stained with crystal violet, quantified, and imaged. **C)** Primary MECs were used for a primary mammosphere assay where the isolated cells were plated in ultra-low attachment plates for 7 days as described previously (N = 3). Then primary mammospheres were harvested and replated for another 7 days under identical incubation conditions to form secondary spheres. Primary and secondary mammosphere formations were recorded. Values ( **A)**, **B)**, **C)**) are presented as the means ± SEM (\**p* < 0.05, \*\**p* < 0.01 as compared to the corresponding untreated control samples). Representative images from each assay are depicted in the right panels. **D)** Mammary tissue sections were prepared from 17-week-old control mice and mice treated with palbociclib. Representative images of IHC analysis are shown with brown staining indicating SOX2-positive cells. Percentages of SOX2-positive cells were graphed as the means ± SEM (\*\**p* < 0.01). **E)** Stemness markers in the mammary tissues (at 17 weeks of age) of mice from control and palbociclib-treated groups were detected using Western blotting. Protein samples from 3 mice in each group are shown.

### 3.8 Palbociclib coordinately suppresses the proliferation- and stemness-associated signaling networks in premalignant mammary tissues

To uncover the underlying mechanism that accounts for the palbociclib induced changes in mammary epithelial cell proliferation and stemness, we examined signaling pathways implicated in ErbB2-driven mammary tumorigenesis, including Cyclin D1-CDK4/6-RB-E2F, ERα-associated, ErbB2-related, and Wnt/β-catenin signaling pathways.

Given that palbociclib directly targets CDK4/6 activity, we firstly examined the Cyclin D1-CDK4/6–RB–E2F cell cycle axis and confirmed the effective inhibition of CDK4/6 signaling in vivo (Figure 8A). Although CDK4 protein levels showed little or only modest reduction, E2F1 and Cyclin D1 were substantially decreased, particularly in the 150 mg/kg group. CDK1 was also reduced, further supporting inhibition of cell cycle progression and the Cyclin D1-CDK4/6-RB-E2F transcriptional program (Figure 8A).

**Figure 8:**
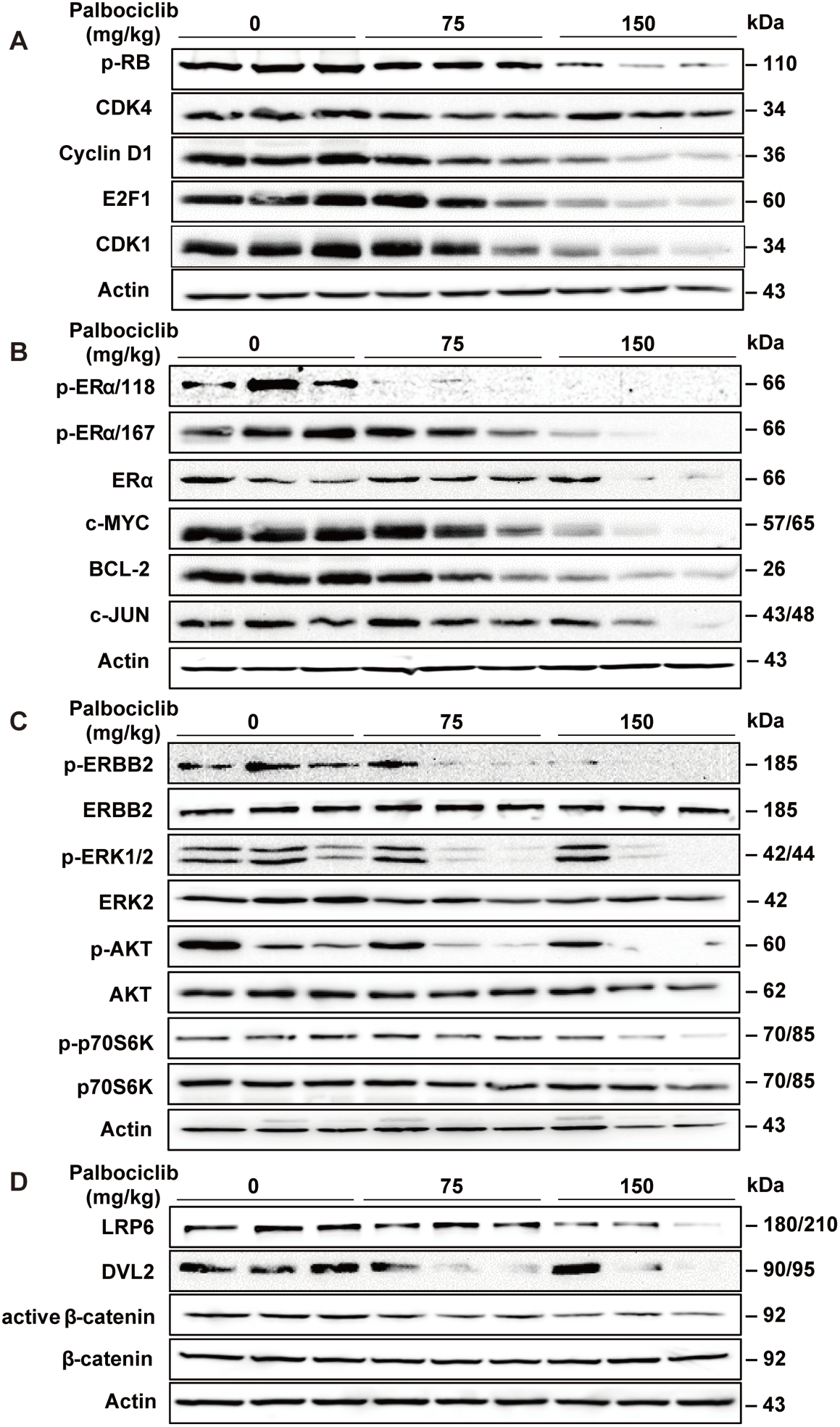
Palbociclib inhibited cell-cycle, ERα-associated, ErbB2-related and Wnt/β-catenin signaling pathways in MMTV-ErbB2 mice. Protein lysate was extracted from mammary tissues of the mice with different treatments (Control, 75 mg/kg and 150 mg/kg every 3 days for 4 weeks), followed by western blot analysis of indicated markers in Cyclin D1-CDK4/6-RB-E2F pathway **A**) ER **B**) ErbB2 **C)** and **D**) Wnt pathways.

Because ERα signaling promotes Cyclin D1 expression and is functionally linked to CDK4/6 activity, we next examined ERα signaling associated molecules. Palbociclib treatment produced a modest decrease in total ERα but a more pronounced reduction in phosphorylated at Ser118 (p-ERα/118) and Ser167 (p-ERα/167) of ERα (Figure 8B). In parallel, the protein expression of several ER-associated downstream molecules, including c-MYC, BCL-2, and c-JUN, was markedly reduced. These findings suggest that palbociclib attenuates ERα-endowed proliferative and survival advantages in premalignant mammary tissues. Given that aberrant RTK/PI3K/AKT signaling is a major downstream effector of ErbB2 and plays an important role in mammary tumorigenesis, we next examined the effects of palbociclib on this pathway. Palbociclib reduced the phosphorylation levels of ERBB2, AKT, and ERK, indicating suppression of RTK-linked proliferative signaling (Figure 8C). Consistently, the phosphorylation levels of mTOR downstream effectors including p70S6K and 4EBP1 were also markedly reduced (Figure 8C), indicating the broad suppression of ErbB2/PI3K/AKT/mTOR signaling. Next, to explore the molecular basis underlying the inhibitory effects of palbociclib on mammary stem/progenitor cells, we examined key components of the Wnt/β-catenin signaling pathway. Palbociclib treatment reduced the protein expression of LRP6 and DVL2 as well as the active β-catenin protein levels in premalignant mammary tissues (Figure 8D).

Collectively, these data indicate that palbociclib exerts broad inhibitory effects across multiple signaling pathways, including Cyclin D1-CDK4/6-RB-E2F, ERα-associated, RTK/PI3K/AKT, and Wnt/β-catenin axes, resulting in attenuation of proliferation and mammary epithelial stemness, ultimately contributing to altered mammary morphogenesis and delayed tumor latency.

## 4. Discussion

In the present study, we found that early and short-term exposure to palbociclib during the premalignant stage can prevent mammary tumorigenesis, which was in parallel with mammary gland growth inhibition in MMTV-ErbB2 transgenic mice. The preventive effect was attributed to the attenuation of MEC subpopulations by reprogramming MaSC differentiation as well as inhibiting epithelial cell proliferation in the premalignant mammary tissues. The underlying mechanisms involve palbociclib coordinately suppresses the complicated proliferation- and stemness-associated signaling networks in premalignant mammary tissues of MMTV-ErbB2 transgenic mice. Collectively, our findings provide proof of concept that short-term CDK4/6 inhibition during an early risk window may become a feasible chemopreventive strategy for ErbB2-driven BC.

CDK4/6 inhibitors are a major class of targeted agents that suppresses RB phosphorylation, thereby inhibiting E2F-mediated transcription and blocking the G1-S transition, ultimately leading to the suppression of cell proliferation [31,32]. Palbociclib was the first CDK4/6 inhibitor to enter clinical practice and has become the prototype of this class of therapeutic agents. Beyond its canonical anti-proliferative activity, accumulating evidence indicates that CDK4/6 inhibition, particularly by palbociclib, regulates multiple biological processes, including cellular senescence, metabolic reprogramming, autophagy, immune surveillance, and epithelial–mesenchymal transition, highlighting the expanded functions of CDK4/6 in cell biology and physiology [33–39]. Clinically, the landmark PALOMA-1, PALOMA-2, and PALOMA-3 trials established palbociclib plus endocrine therapy as a standard treatment for HR-positive/ErbB2-negative advanced BC and showed significant benefits in progression free survival [40–43]. Although initially approved for HR-positive/ErbB2-negative BC, accumulating evidence suggests that palbociclib also has the therapeutic value in ErbB2-positive neoplasm [44–47]. More recently, phase III PATINA trial demonstrated that addition of palbociclib to maintenance anti-ErbB2 and endocrine therapy significantly improved progression free survival (44.3 vs. 29.1 months; HR = 0.75, 95% CI: 0.59–0.96; *p* = 0.02) in patients with HR-positive/ErbB2-positive metastatic BC [44]. In the case of CDK4/6 inhibition in ErbB2-driven BC model, numerous preclinical studies showed that palbociclib suppresses the cell proliferation of ErbB2-positive BC cells and synergizes with ErbB2-targeted therapies by inhibiting the downstream Cyclin D1-CDK4/6-RB axis [47]. Consistent with these findings, our data also demonstrated that palbociclib inhibits in vivo cell proliferation and in vitro allograft tumor growth in the context of ErbB2 overexpression. However, most of the existing studies have mainly focused on established tumors, but the impact of palbociclib on ErbB2-driven tumorigenesis at the premalignant stage remains largely unexplored.

Genetic ablation of Cyclins D1, CDK4, or CDK6 exhibits a decreased predisposition to tumorigenesis in a variety of neoplasms [48]. For instance, CCND1- or CDK4-null mice, or knock-in mice expressing kinase-inactive cyclin D1-CDK4/6, were resistant to develop ErbB2-driven BC [10–12,14]. These observations indicated that CDK4 and CDK6 might represent excellent targets in cancer prevention. Although accumulating preclinical evidence supports the potential of palbociclib as a chemopreventive agent for individuals at high risk of ErbB2-positive BC, the optimal timing, duration, and treatment regimen for clinical prevention remain to be established. To address this question, we employed the MMTV-ErbB2 transgenic mouse model, a well-established spontaneous model that faithfully recapitulates the key features of human ErbB2-driven BC, including the progression from premalignant hyperplasia to invasive carcinoma [49,50]. This model has been widely used to evaluate chemopreventive strategies, including rexinoids, lapatinib, buformin, providing a robust in vivo platform for assessing preventive interventions within a common ErbB2-driven biological context [17,51,52].

In the present study, palbociclib was administered for 8 weeks (100 mg/kg, every 3 days) during the premalignant risk window before tumor onset. We demonstrated that this treatment regimen significantly delayed the initial onset of mammary tumor by 9 weeks in MMTV-ErbB2 transgenic mice, and even completely prevented tumor formationin a subset of animals (Figure 4B). Specifically, the median tumor latency was prolonged from 38 weeks in the control group to 55 weeks in the palbociclib-treated group following only 8 weeks of treatment. Consistent with our hypothesis, these findings demonstrate that short-term palbociclib exposure prior to tumor initiation induced long-lasting cancer preventative effects. Consistent with our findings, only 4 weeks of daily palbociclib treatment reduced liver tumor incidence from 100% to 29.15% in using Fah-/- mice and N-Ras+ AKT mouse model of hepatocellular carcinoma, accompanied by significant decreases in both tumor number and tumor size [53]. Notably, the reduction in tumor incidence was more pronounced in the liver cancer models (approximately 71%) than in our study, in which 3 of 15 mice (20%) remained tumor-free. This difference may partly ascribe to the different treatment regimens: palbociclib was administered each day for 28 days in the liver cancer study, whereas mice in our study received a total of 18 doses every 3 days. Moreover, differences in tumor type and the underlying mechanisms of tumorigenesis may also contribute to the different preventive responses. In the clinical setting, prolonged using of palbociclib suppressed immunocompetence, necessitating a one-week break after every three weeks of oral administration [54]. More similar to our treatment schedule, but direct extrapolation of the doses used in mice to human-equivalent doses should be interpreted cautiously because of interspecies differences in pharmacokinetics, drug exposure, and tolerability.

Reduced mammary tumor risk was accompanied by the modified morphogenesis that showed the decreased MEC density and proliferative index in the premalignant mammary tissues [26,55] (Figure 5). Based on these data, outstanding questions remain regarding the molecular and cellular mechanisms that result in longer tumor latency and remarkable morphogenic changes in the premalignant tissues.

Recent progress in stem cell research indicates that mammary stem cells contribute to tumor heterogeneity, initiation, recurrence, and invasive potential through the differentiation into cancer stem cells (CSCs)[29,30,55–59]. Therefore, studying the regulation of MaSCs/progenitor cells in mammary development is critical for understanding breast cancer etiology and associated risk factors. In this context, we focused on palbociclib inhibition of MEC stemness in context with modified tumorigenesis. Our data showed that palbociclib induced marked changes in MEC subpopulations that comprised the significantly reduced the luminal subpopulation (CD24^high^CD49f^low^) and MRU enriched subpopulation (CD24^high^CD49f^high^) (Figure 6). Because CD24highCD49f^low^ MECs constitute the major epithelial cell subpopulation of mammary ducts and alveoli [20,60], their reduction may explain the decreased MEC density and diminished alveolar budding observed in the mammary gland whole mounts after palbociclib exposure (Figure 5). The reduction in the MRU enriched population further suggests that palbociclib suppresses MaSCs self-renewal. Consistent with these phenotypic changes, CFC, mammosphere, and 3D culture assays further demonstrated that palbociclib reduced the abundance of luminal progenitor cells and impaired the self-renewal capacity of MaSCs in premalignant mammary tissues (Figure 7).

Previous studies have shown that the CD61^high^CD49f^mid^ MEC population is enriched for luminal progenitor cells and share a similar phenotype with TICs, which are considered candidate cells of origin for ErbB2-driven mammary tumorigenesis [17,19,56]. In the present study, palbociclib significantly reduced the CD61^high^CD49f^mid^ MEC subpopulation (Figure 6), suggesting that short-term CDK4/6 inhibition restricts the expansion of TICs. This effect may reduce the pool of cells susceptible to oncogenic transformation, thereby delaying ErbB2-driven mammary tumor initiation and progression. Importantly, as we show in our studies, the selective targeting of CSCs/ TICs, may contribute to the preventive effects of palbociclib to block pro-oncogenic events that cause cancer initiation in premalignant tissues and the progression of cancer at various stages. Understanding theunderlying mechanisms that result in selective inhibition of CSC/TIC populations is therefore important for explaining its potential preventive effects. As a potentially critical pathway for the CSCs targeted effects of palbociclib, Wnt/β-catenin pathway plays a substantial role in the regulation of numerous pro-cancerous cellular responses, including cell differentiation and proliferation [61–65]. Our data showed that palbociclib inhibited β-catenin activation and reduced stemness-associated markers (NANOG, KLF4, SOX2 and OCT4), providing a potential connection between the differential epithelial subpopulations following short-term palbociclib exposure in premalignant mammary tissues (Figure 6-7). Due to the concurrent reduction in the CD61^high^CD49f^mid^ cell subpopulation and mammosphere formation efficiency in the MECs from palbociclib exposure mice, our results suggest that palbociclib may induce Wnt/β-catenin-mediated MaSC reprogramming and deter the differentiation into CSCs. This proposed mechanism would explain the selective targeting of CSCs by metformin and buformin, as previously published in our lab [17,66].

In addition to the inhibitory effects on cell stemness, palbociclib also displayed substantial inhibition of epithelial cell proliferation from preneoplastic mammary glands of MMTV-ErbB2 mice. Previously, we reported that receptor tyrosine kinases inhibitor Lapatinib, reduce growth inhibitory and ultimate tumor prevention through the suppression of ErbB2 and ERα signaling pathways [52]. Recent reports further show that extensive crosstalk exists between ERα signaling and the cell-cycle regulatory network. Estrogens bind the nuclear ERα, inducing binding at genomic estrogen response elements and subsequent gene transcription. One of the gene products is Cyclin D1, which binds CDK4/6 to drive progression through the cell cycle. Cyclin D1 associates with CDK4/6 to activate the CDK4/6-RB-E2F signaling axis, thereby promoting G1/S cell-cycle progression [67–69]. In addition, Cyclin D1 directly enhances ERα transcriptional activity, forming a positive feedback loop that promotes cell proliferation independently of estrogen [70,71]. Our findings suggest that palbociclib disrupts this positive feedback loop, resulting in coordinated suppression of both CDK4/6-RB-E2F signaling and ERα signaling, thereby inhibiting cell proliferation. Moreover, those results also suggest that palbociclib may induce a broader impact on positive feedback loop between cell cycle and ERα signaling beyond the canonical regulatory pathways. For instance, concurrent inhibition of ErbB2 and ERα in mammary tissues suggests that palbociclib may block the crosstalk between these two signaling molecules, leading to downstream effects on multiple pathways. Similarly, metformin and buformin, two drugs for diabetes, also induced inhibition of ERα-ErbB2 crosstalk and the consequent cellular responses are likely contributors to cell proliferation and mammary tumor development [17,66]. Beyond the suppression of proliferation and stem/progenitor-associated properties observed in our study, CDK4/6 inhibition may exert preventive effects through additional mechanisms, such as inducement of senescence that warrants further investigation.

We acknowledge that palbociclib induced broad suppression of multiple signaling pathways in premalignant mammary tissues; however, the underlying molecular mechanisms remain incompletely understood. To address this, transcriptomic profiling by RNA sequencing is currently underway in our laboratory. This analysis will provide a comprehensive view of the transcriptional and signaling networks regulated by short-term CDK4/6 inhibition during the premalignant risk window and may identify additional molecular targets and biomarkers for evaluating chemopreventive efficacy. In addition, CDK4/6 inhibition has been reported to remodel the tumor immune microenvironment by enhancing antitumor immunity through upregulation of MHC class I expression and selective depletion of regulatory T cells [38] [39]. Although the present study focused primarily on mammary epithelial cells, it is possible that palbociclib also reprograms the immune microenvironment of premalignant mammary tissues, thereby contributing to its long-lasting chemopreventive effects. This hypothesis warrants further investigation. From a translational perspective, future studies are required to optimize the timing and dosing of preventive CDK4/6 inhibition, identify biomarkers for stratifying women at high risk of ErbB2-positive BC, and evaluate whether short-term CDK4/6 inhibitor exposure can provide durable clinical benefit. Furthermore, combining CDK4/6 inhibitors with anti-ErbB2 therapy, endocrine therapy, or other targeted agents may further enhance chemopreventive efficacy and broaden their applicability across different BC subtypes.

## 5. Conclusion

In conclusion, our study demonstrates that short-term palbociclib exposure during the early premalignant risk window suppresses mammary tumorigenesis in MMTV-ErbB2 transgenic mice through MaSC reprogramming and inhibition of mammary stem/progenitor cell stemness. Luminal progenitor cells and the MRU subpopulation stemmed from MaSCs are likely the targets of palbociclib, which may contribute to the observed delay in mammary tumor development. Mechanistically, palbociclib inhibited multiple interconnected signaling networks including CDK4/6-RB-E2F, ER-associated, ErbB2-related and Wnt/β-catenin signaling pathways, all of which synergistically reduces the risk of mammary tumorigenesis. These findings provide a mechanistic foundation for CDK4/6 inhibitor-based BC prevention and early term intervention, and support further preclinical and clinical investigation to extend the use of CDK4/6 inhibitors from the treatment of established to the premalignant setting.

## 6. Patents

This section is not mandatory but may be added if there are patents resulting from the work reported in this manuscript.

## Author Contributions

**Y.L.**: Data curation and analysis, writing – review & editing; **A.B.P.**: Data curation and analysis, review & editing; **Z.M.**: Data curation and analysis; **L.W.**; Data analysis and interpretation; **Y.Q.**: Data analysis, writing-drafting, review & editing; **X.Y.** Conceptualization, funding acquisition, investigation, project administration, writing – drafting, editing & review. All authors have read and agreed to the published version of the manuscript.

## Funding

This work was supported in part by a R16 grant from the National Institute of General Medical Sciences (1R16GM145545) to X.Y., a U54 grant from the National Institute on Alcohol Abuse and Alcoholism (U54 AA019765), and a RCMI U54 grant from the National Institute on Minority Health and Health Disparities (U54 MD012392).

## Institutional Review Board Statement

The animal study protocol was approved by the Institutional Animal Care and Use Committee of the North Carolina Research Campus (protocol code 23-003 and date of approval 4/17/2023).

## Informed Consent Statement

Not applicable.

## Data Availability Statement

All data is contained within the manuscript.

## Conflicts of Interest

The authors declare no conflicts of interest.

## Abbreviations

The following abbreviations are used in this manuscript

ErbB2/HER2: Human epidermal growth factor receptor 2
MEC: Mammary epithelial cell
MRU: Mammary reconstitution unit
TIC: Tumor-initiating cell
BC: Breast cancer
MaSC: Mammary stem cell
CDK4/6: Cyclin-dependent kinase 4/6
IHC: Immunohistochemistry
ECL: Enhanced chemiluminescence
CFC: Colony-Forming cell
SEM: Standard error
Myo: Basal/myoepithelial cells
CSC: Cancer stem cell
CCK-8: Cell Counting Kit-8

## Notes

### Competing Interest Statement

The authors have declared no competing interest.

